# Response inhibition and error-monitoring in cystinosis (CTNS gene mutations): Behavioral and electrophysiological evidence of a diverse set of difficulties

**DOI:** 10.1101/2023.03.31.535145

**Authors:** Ana A. Francisco, John J. Foxe, Alaina Berruti, Douwe J. Horsthuis, Sophie Molholm

**Affiliations:** The Cognitive Neurophysiology Laboratory, Department of Pediatrics, Albert Einstein College of Medicine, Bronx, New York, USA; Department of Neuroscience, Rose F. Kennedy Center, Albert Einstein College of Medicine, Bronx, New York, USA; The Frederick J. and Marion A. Schindler Cognitive Neurophysiology Laboratory, Ernest J. Del Monte Institute for Neuroscience & Department of Neuroscience, University of Rochester School of Medicine and Dentistry, Rochester, New York, USA

**Keywords:** Go/No-Go, EEG, N2, P3, Pe

## Abstract

Cystinosis, a rare lysosomal storage disease, is characterized by cystine crystallization and accumulation within tissues and organs, including the kidneys and brain. Its impact on neural function appears mild relative to its effects on other organs, but therapeutic advances have led to substantially increased life expectancy, necessitating deeper understanding of its impact on neurocognitive function. Behaviorally, some deficits in executive function have been noted in this population, but the underlying neural processes are not understood. Using standardized cognitive assessments and a Go/No-Go response inhibition task in conjunction with high-density electrophysiological recordings (EEG), we sought to investigate the behavioral and neural dynamics of inhibition of a prepotent response and of error monitoring (critical components of executive function) in individuals with cystinosis, when compared to age-matched controls. Thirty-seven individuals diagnosed with cystinosis (7-36 years old, 24 women) and 45 age-matched controls (27 women) participated in this study. Analyses focused on N2 and P3 No-Go responses and error-related positivity (Pe). Atypical inhibitory processing was shown behaviorally. Electrophysiological differences were additionally found between the groups, with individuals with cystinosis showing larger No-Go P3s. Error-monitoring was likewise different between the groups, with those with cystinosis showing reduced Pe amplitudes.

## INTRODUCTION

Cystinosis, with an estimated incidence of 1 per 100,000 to 200,00 live births (1), is an autosomal recessive lysosomal storage disease caused by bi-allelic mutations in the 17p13.2-located CTNS gene (2). These mutations, more common in populations of Northern Europe and North America of European descent (1, 3), result in dysfunctional transport of cystine and, ultimately, in cystine accumulation and crystal formation in various tissues and organs. Cystinosis, in which this dysfunction is systemic, has been associated with renal, retinal, endocrinological, muscular, and neurological complications (4, 5).

Subcortical and cortical atrophy, Chiari I malformation, white matter abnormalities, and atypical electrophysiological (EEG) activity are among the neurological findings described in individuals with cystinosis (6–12). These findings may account for some of the differences in cognitive function and academic performance reported in this population (13–22). The neurocognitive profile associated with CTNS mutations and its developmental path is, nevertheless, still poorly understood.

Executive functioning, which refers to a set of high-level cognitive processes underlying goal-directed behaviors, is of particular interest given that several of its components—such as memory updating, task/attention shifting, goal monitoring, and inhibition of inappropriate responses—are critical for and predictive of academic and professional achievements (23–27), as they optimize approaches to unfamiliar circumstances and contribute to performance monitoring and self-regulation (28–31).

The small number of behavioral studies that investigated executive function in cystinosis suggest the presence of some difficulties in this population (14, 32, 33), but not consistently. For instance, while Ballantyne and colleagues showed worse performance across different components of executive functioning in those with cystinosis (32), Besouw et al observed differences between individuals with and without cystinosis only in sustained attention, not in other executive functioning components (14). Our own work suggests no (behavioral) verbal working memory difficulties in a small group of adults living with cystinosis (34), but some differences between individuals with cystinosis and age-matched controls in the amplitude of event-related potentials (ERPs) associated with sensory memory and attentional processes (34, 35).

In the current study, we focused on two components of executive function: response inhibition and error monitoring. Response inhibition is the process by which one suppresses a prepotent response that might be irrelevant or inappropriate in a given context and is essential for adjusting behavior dynamically with changing environmental contexts (36–38). Previous research suggests maintained response inhibition in some children with cystinosis, who completed a standardized inhibition task (14). Error monitoring relates to the identification and correction of deviance from a correct response (39) and is required to achieve goal-directed behavior and maintain task performance (40). To our knowledge, error monitoring has not been investigated in cystinosis.

To assess response inhibition and error monitoring, we used standardized cognitive measures and a Go/No-Go EEG task and compared behavioral and neural responses of those with cystinosis to those of an age-matched control group. The analyses focused on reaction-time and d’ measures of response inhibition and on event-related potential components typically evoked during similar Go/No-Go tasks: The No-Go N2, a negative-going ERP component peaking between 200 and 300 ms and representing early, automatic inhibitory (41–44) and/or conflict detection processes (45–47); the No-Go P3, a positive potential that peaks at about 300-500 ms, argued as a marker of response inhibition (48–52), stimulus evaluation (53–55) and adaptive, more effortful forms of control (43, 44, 56); and the error-related positivity (Pe), a component peaking between 200 and 500 ms post incorrect-response, which has been suggested to reflect conscious error processing or updating of error context (57, 58). Lastly, we measured the relationship between brain responses and cognitive function.

Though our previous work does not suggest extensive differences in cognitive and neural function in cystinosis, there is anecdotal evidence of somewhat pervasive difficulties in executive function type abilities in this population. Hence, slower and/or less accurate responses and reduced N2/P3 and Pe amplitudes might be observed in individuals living with cystinosis when compared to their age-matched peers. A better characterization of the cognitive profile associated with cystinosis is critical to developing effective interventions to compensate for or improve areas of cognitive vulnerability.

## MATERIALS AND METHODS

### Participants

Thirty-eight individuals diagnosed with cystinosis (CYS; age range: 7-36 years old, 24 women) and 45 age-matched controls (CT; 27 women) were recruited. Those with cystinosis were primarily recruited via family and foundation groups in social media channels, while controls were recruited via flyers in the surrounding neighborhood and a lab-maintained participant database. Exclusionary criteria for the control group included developmental and/or educational difficulties or delays, neurological problems, and severe mental illness diagnosis. Exclusionary criteria for individuals with cystinosis included current neurological problems. All participants had normal or corrected to normal vision. A Snellen chart was used to assess visual acuity and participants were asked at the start of the EEG paradigm if they could easily see the stimuli and their different components. Due to illness on the scheduled day of testing, one individual with cystinosis was excluded from the final sample. All individuals, or their legal guardian when under 18 years old, signed a consent form. Participants were monetarily compensated for their time. This study and all the associated procedures were approved by the Albert Einstein College of Medicine Institutional Review Board. All aspects of the research conformed to the tenets of the Declaration of Helsinki.

### Experimental Procedure and Stimuli

Participation consisted of two visits, which involved completion of a cognitive function battery and EEG recordings. For the cognitive battery, verbal and non-verbal intelligence was assessed using age-appropriate Wechsler Intelligence Scales. To assess response inhibition, the Conners Continuous Performance Test 3, CPT (59), and the Color-Word Interference Test of the Delis-Kaplan Executive Function System, D-KEFS (60) were used. The CPT is a brief visual inhibition task which indexes various measures of attention, including inattentiveness, impulsivity, sustained attention, and vigilance. Here, we focused on Commissions (i.e., incorrect response to non-target stimuli) and Perseverations (i.e., anticipatory responses). Due to software malfunctioning, seven individuals with cystinosis and six controls did not complete the CPT and were therefore not included in the related analyses.

The Color-Word Interference Test of the D-KEFS consists of four tasks, including color naming, word reading, inhibition, and inhibition/switching. As with the CPT, we focused on measures most relevant to response inhibition, and therefore, only included the inhibition score in our analyses. The inhibition score was calculated using the inverse efficiency score (*IES=RT/(1-PE)*, where *RT* is the individual’s average reaction time in the condition, and *PE* is the subject’s proportion of errors in the condition (61). Four controls and two individuals with cystinosis did not complete their second testing session and thus the Color-Word Interference Test and were therefore not included in the related analyses.

During the EEG recording session, participants were asked to respond as quickly and as accurately as possible to a visual response-inhibition task. Positive and neutral valence images from the International Affective Picture System (IAPS; Lang and Cuthbert, 1997) were presented in a pseudorandom sequence. Participants were instructed to press the left mouse button upon each stimulus presentation, unless the stimulus was a repetition of the immediately preceding stimulus, in which case they should withhold (inhibit) their response. Stimuli, subtended 8.6° horizontally by 6.5° vertically, were presented centrally every 1000 ms for 600 ms with a (random) inter-stimulus-interval between 350 and 450ms (Figure 1). Three 12-minute blocks were run. Each block consisted of 540 trials, for a total of 1620 trials per participant, 243 of which were inhibition trials.

**Figure 1.**
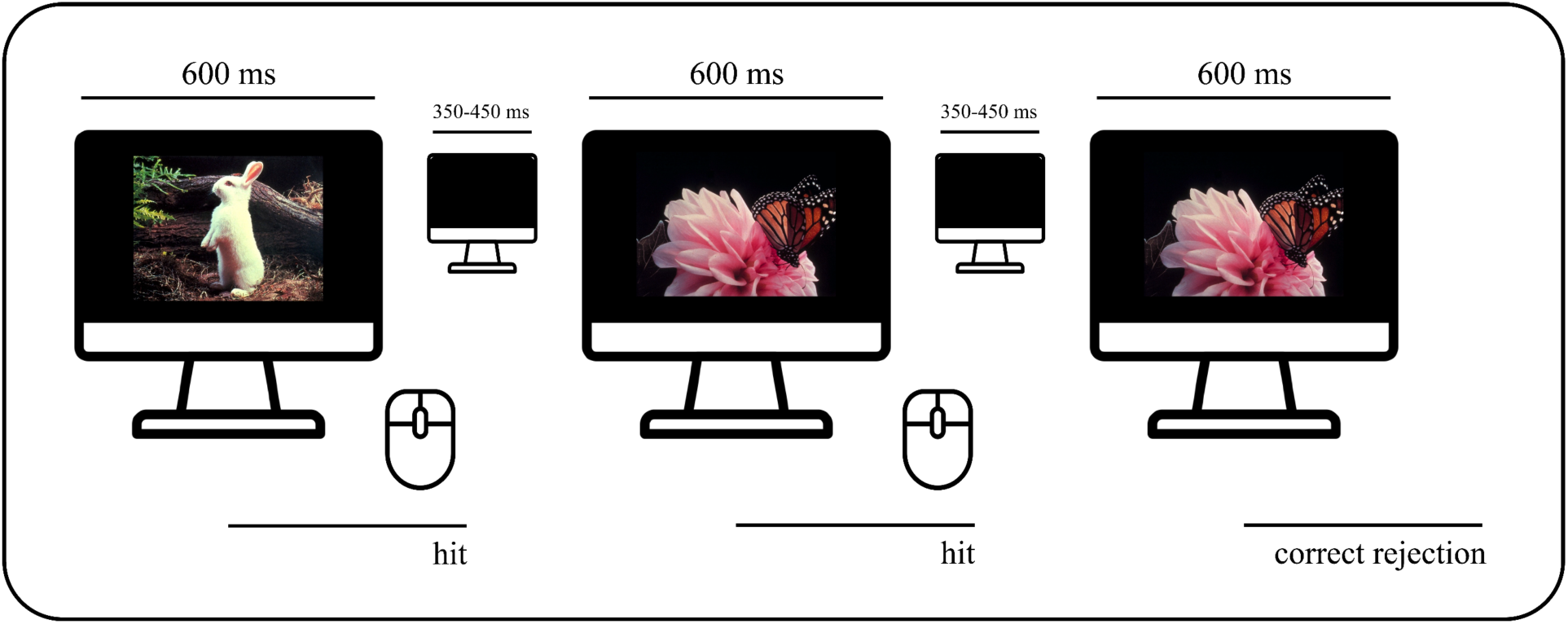
Go/No-Go EEG task.

### Data acquisition and analysis

Continuous EEG data were recorded from 64 scalp electrodes at a sampling rate of 512 Hz (Active 2 system; Biosemi^tm^, The Netherlands; 10-20 montage) and then preprocessed using the EEGLAB toolbox (version 2021.0) (62) for MATLAB (version 2021a; MathWorks, Natick, MA) (the full pipeline can be accessed at: https://github.com/DouweHorsthuis/EEG_to_ERP_pipeline_stats_R) (63). Preprocessing steps included down-sampling data to 256 Hz, re-referencing to the average, and filtering with a 0.1 Hz high pass filter (0.1 Hz transition bandwidth, filter order 16896) and a 45 Hz low pass filter (11 Hz transition bandwidth, filter order 152). Both were zero-phase Hamming windowed sinc FIR filters. Noisy channels were excluded based on kurtosis and visual confirmation. Artifacts from blinks and saccades were eliminated via Independent Component Analysis (ICA). The spherical spline method was then used to interpolate channels that were removed in previous steps. Data were segmented into epochs of -100 ms to 1000 ms using a baseline of -100 ms to 0 ms. For the error-related activity analyses, data were segmented into response-locked epochs of -100 ms to 700 ms using a baseline of -100 ms to 0 ms. All epochs went through an artifact detection algorithm (moving window peak-to-peak threshold at 120 µV). To equate number of trials per participant, 200 trials for hits, 50 trials for correct rejections, and 50 trials for false alarms were chosen randomly per subject.

### Response inhibition related ERPs

Time windows and electrode locations were selected based on past research and confirmed (and adjusted) by inspecting grand averages collapsed across the groups. N2 was measured between 210 and 240 ms at AFz and Fz and P3 between 350 and 500 ms at FCz and Cz (64–66). Error-related positivity (Pe) was measured between 200 and 400 ms at CPz (67, 68). Mean amplitude data were used for both between-groups statistics and Spearman correlations. Behavioral measures (accuracy and reaction time) were additionally taken during the EEG task. Hits were defined as responses to a non-repeated picture; correct rejections as the absence of response to a repeated picture; false alarms as responses to a repeated picture. Only hits and correct rejections preceded by a hit were included. *D*-prime (d’ = z(H) - z(F)) was calculated per subject. All *p-values* (from *t*-tests and Spearman correlations) were submitted to Holm-Bonferroni corrections for multiple comparisons (69), using the *p.adjust* of the *stats* package in R (70). Mixed-effects models were implemented to analyze trial-by-trial data, using the *lmer* function in the *lme4* package (71) in R (70). Group was always a fixed factor, and trial type an additional numeric fixed factor for reaction times and N2/P3 analyses. Subjects and trials were added as random factors. Models were fit using the maximum likelihood criterion. *P* values were estimated using *Satterthwaite* approximations.

## RESULTS

### Demographics and cognitive function measures

Table 1 shows a summary of the included participants’ age, sex, and cognitive functioning (verbal IQ and perceptual reasoning and inhibition-related measures: D-KEFS color-word inhibition composite measure and CPT commissions and perseverations). Two-sample independent-means *t* tests were run in R (70) to test for group differences in age and cognitive performance. In cases in which the assumption of the homogeneity of variances was violated, *Welch* corrections were applied to adjust the degrees of freedom. A chi-square test was run to test for independence between sex and group. Effect sizes were calculated utilizing Cohen’s *d* and *w*.

**Table 1.** Characterization of the control and cystinosis individuals included in the analyses: Demographics and cognitive function (IQ and inhibition measures).

|  | control | cystinosis | statistical test | effect sizes |
| --- | --- | --- | --- | --- |
| **age** | M=17.36; SD=8.92 | M=17.62; SD=9.36 | $t=-0.13, df=75.43, p=.90$ | $d=0.03$ |
| **sex** | 27 F, 18 M | 25 F, 12 M | $\chi^2=0.23, df=1, p=.63$ | $w=0.08$ |
| **verbal IQ** | M=109.71; SD=19.85 | M=93.46; SD=11.21 | $t=4.66, df=71.57, p<.01$ | $d=1.00$ |
| **perceptual reasoning** | M=103.44; SD=13.49 | M=85.19; SD=13.70 | $t=6.04, df=76.50, p<.01$ | $d=1.36$ |
| **inhibition composite (d-kefs)\*** | M=61.19; SD=24.63 | M=83.08; SD=40.39 | $t=-2.56, df=57.44, p<.05$ | $d=0.62$ |
| **commissions (cpt)\*** | M=43.83; SD=7.29 | M=55.93; SD=9.77 | $t=-4.98, df=53.66, p<.01$ | $d=1.27$ |
| **perseverations (cpt)\*** | M=48.42; SD=5.01 | M=52.82; SD=8.57 | $t=-2.27, df=44.73, p=.06$ | $d=0.60$ |

\* higher scores represent worse performance

As can be seen in Table 1 and Figure 1, the groups differed in verbal and non-verbal abilities, and in inhibition (composite measure and commissions), with individuals with cystinosis performing worse than their age-matched peers. No differences were found in age, sex, or perseverations.

**Figure 1.**
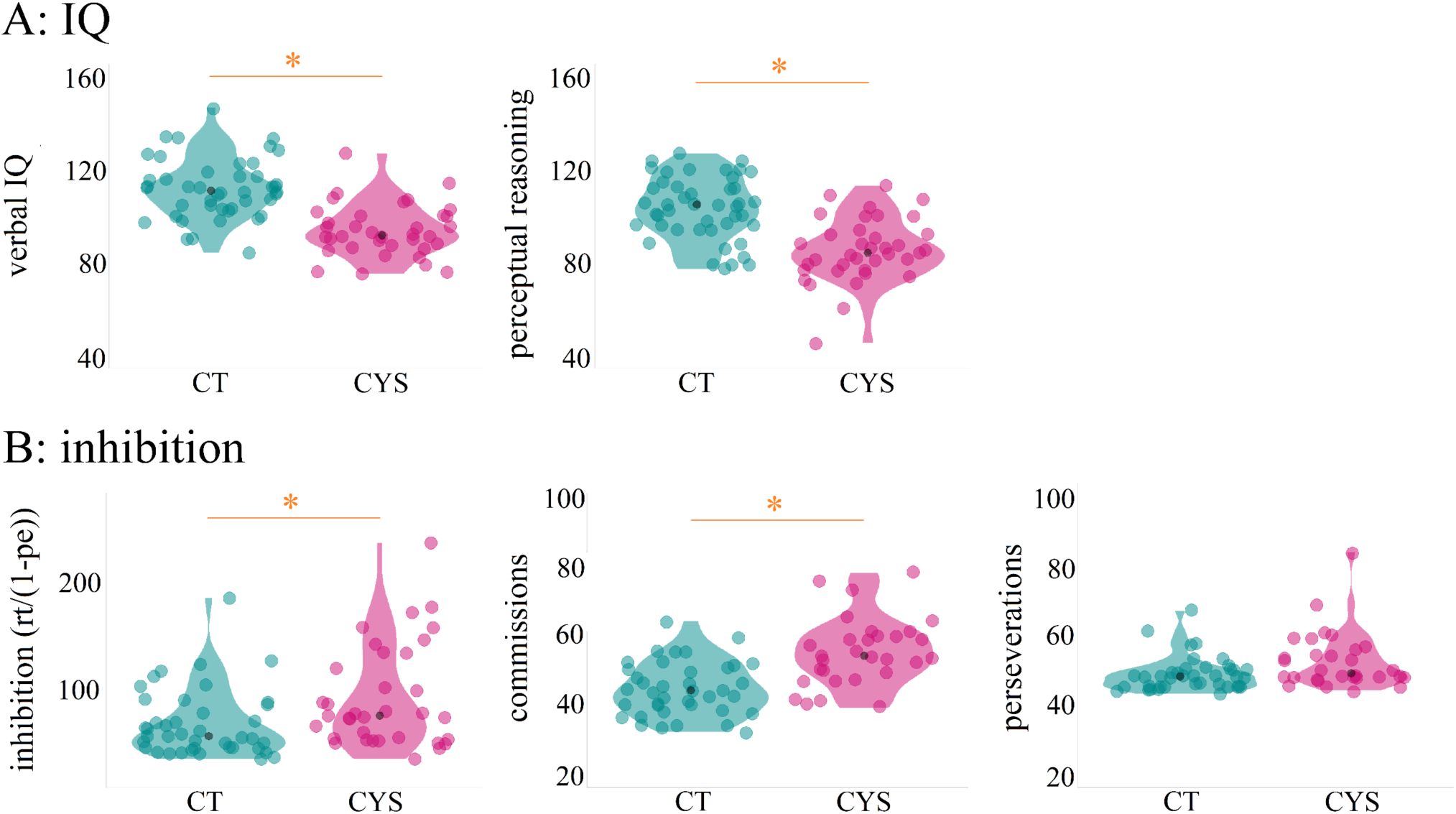
Included participants’ performance on IQ (panel A) and inhibition measures (panel B). In panel B, higher scores represent worse scores.

### Go/No-Go EEG task

#### Behavioral performance

Figure 2 and Table S1 (supplementary materials) show the participants’ behavioral performance (*d-*prime, proportions of hits and false alarms, and reaction times) on the Go/No-Go EEG task. To test for differences in *d-*prime between the groups, two-sample independent-means *t* tests were run in R (70), as described above. When compared to their controls, individuals with cystinosis presented lower *d-*prime scores (*t*=2.70, *df*=70.29, *p*=.01, *d*=0.62), reflecting both lower rates of hits (*t*=2.60, *df*=73.80, *p*=.02, *d*=0.59) and higher rates of false alarms (*t*=-2.09, *df*=62.63, *p*=.04, *d*=0.49) (Figure 2A).

**Figure 2.**
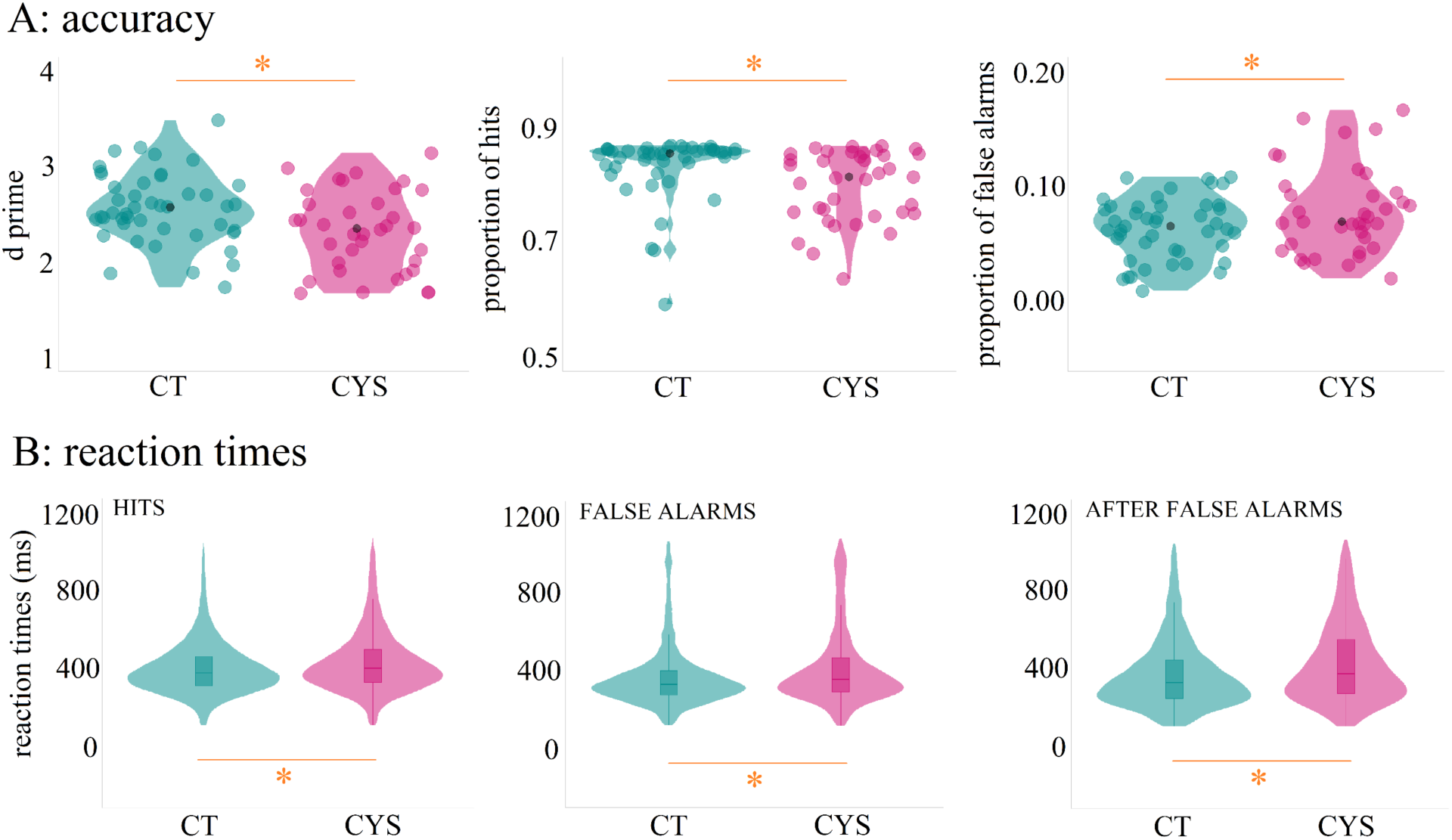
Participants’ behavioral performance on the Go/No-Go EEG task.

To test for differences in reaction time between the groups, mixed-effects models were implemented as described in the Methods Section. There was an overall effect of group: Individuals with cystinosis responded slower than their peers (*ß* = 37.08, SE = 15.53, *p* = .02) (Figure 2 B). There was also an effect of trial type, with false alarms (*ß* = -42.88, SE = 2.38, *p* = .01) and trials after false alarms (*ß* = - 33.12, SE = 2.46, *p* = .01) resulting in shorter reaction times than hits. Additionally, both groups slowed their responses on trials after false alarms, when compared to false alarms (*ß* = 9.38, SE = 4.03, *p* = .02).

#### Response inhibition: N2 & P3

Figures 3 and 4 show the averaged ERPs for N2 (Figure 3) and P3 (Figure 4) by group and trial type (hits and correct rejections). Mixed-effects models were implemented as described in the Methods Section.

**Figure 3.**
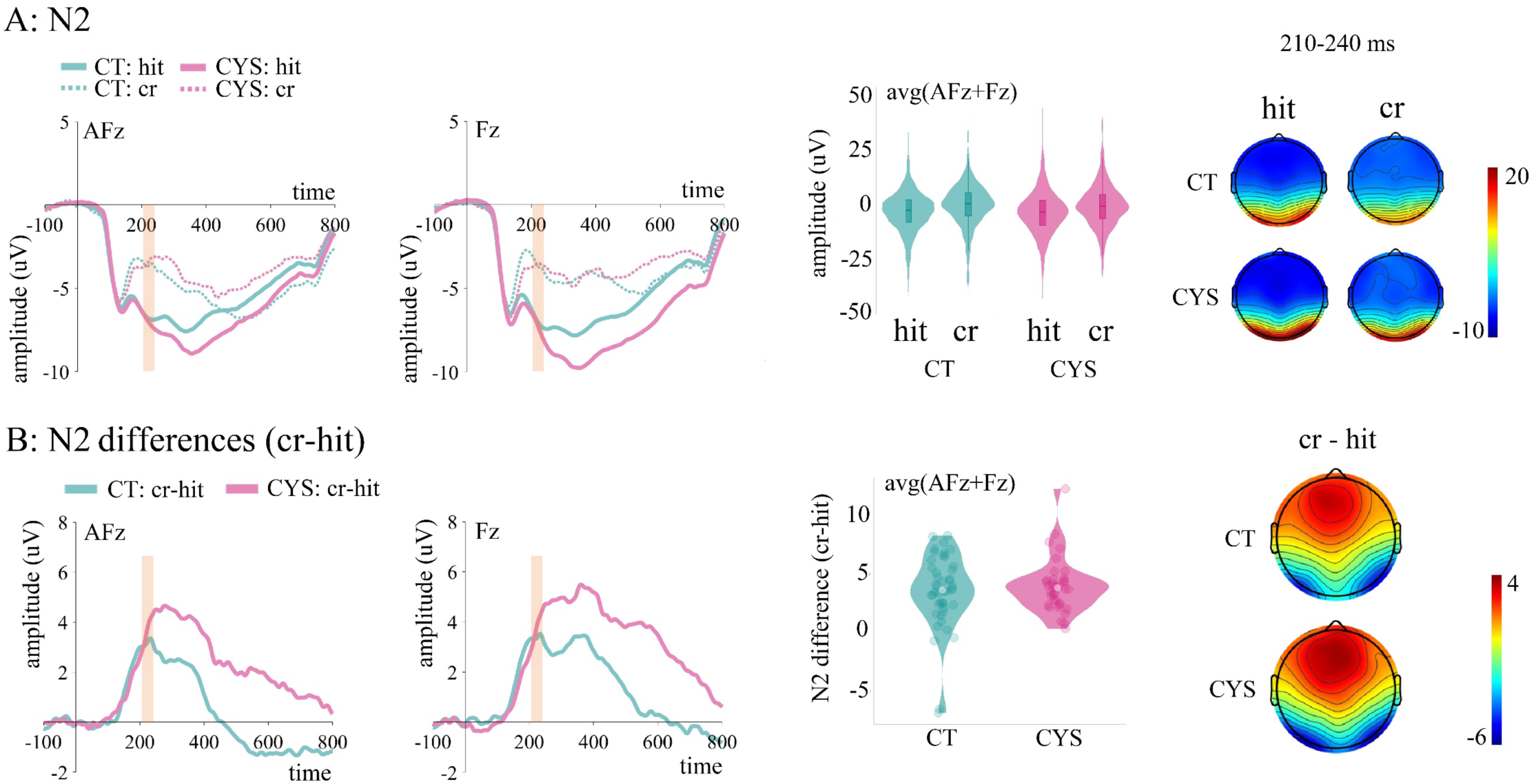
Panel A: Averaged ERPs per group at AFz and Fz, and plots showing distribution of amplitudes for hits and correct rejections per group (trial-by-trial data; average of AFz and Fz) for N2. Panel B: Difference waves (correct rejections – hits) per group, and plots showing distribution of amplitudes for those differences per group at AFz and Fz (averages). Shading in ERP plots indicates time window of interest.

**Figure 4.**
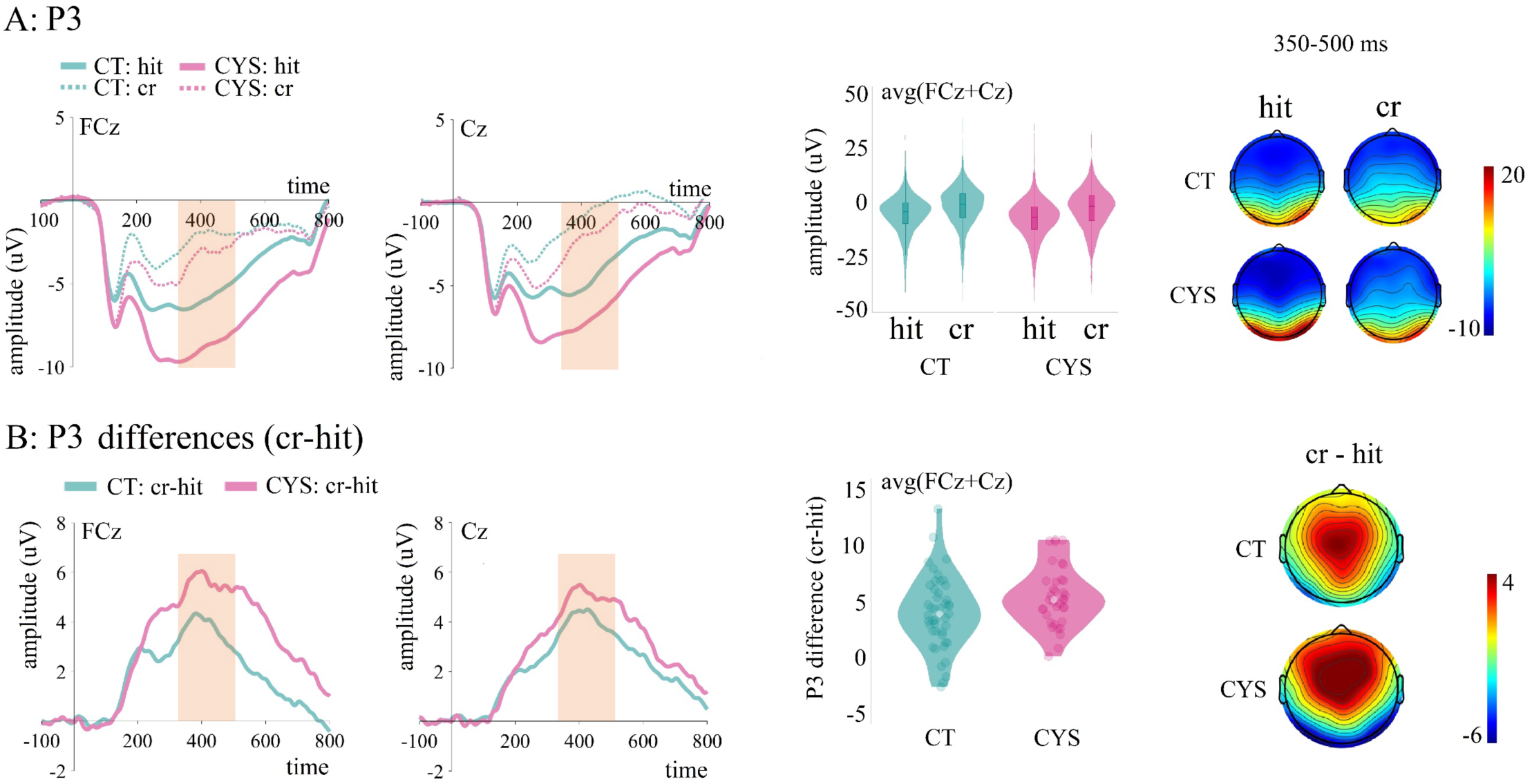
Panel A: Averaged ERPs per group at AFz and Fz, and plots showing distribution of amplitudes for hits and correct rejections per group (trial-by-trial data; average of AFz and Fz) for P3. Panel B: Difference waves (correct rejections – hits) per group, and plots showing distribution of amplitudes for those differences per group at AFz and Fz (averages). Shading in ERP plots indicates time window of interest.

In the N2 time window, no differences were found between the groups (*ß* = -0.36, SE = 0.83, *p* =.66). Both groups presented more positive amplitudes for correct rejections when compared to hits (*ß* = 2.83, SE = 0.30, *p* = .01) (Figure 3A). As can be seen in Figure 3B, cystinosis and control groups presented similar difference amplitudes (correct rejections – hits). In line with this observation, t-tests failed to reveal statistical differences between the groups (*t*=-0.78, *df*=78.25, *p*=.44, *d*=0.17).

As can be appreciated in Figure 4A, individuals with cystinosis presented more negative amplitudes in the P3 time window when compared to their age-matched controls (*ß* = -2.54, SE = 0.99, *p* =.01). Both groups presented more positive amplitudes for correct rejections when compared to hits (*ß* = 3.77, SE = 0.28, *p* = .01) (Figure 4A). Figures 4A and B suggest a larger difference between hits and correct rejections in the cystinosis group. T-tests confirmed modest, but significant differences between the groups (*t*=-2.11, *df*=78.98, *p*=.04, *d*=0.46).

#### Error-related activity: Pe

Figure 5 shows the averaged ERPs for false alarm trials, after incorrect button press. Mixed-effects models, implemented as described in the Methods Section, revealed that when compared to controls, those with cystinosis showed decreased Pe amplitudes (*ß* = -1.62, SE = 0.81, *p* = .04).

**Figure 5.**
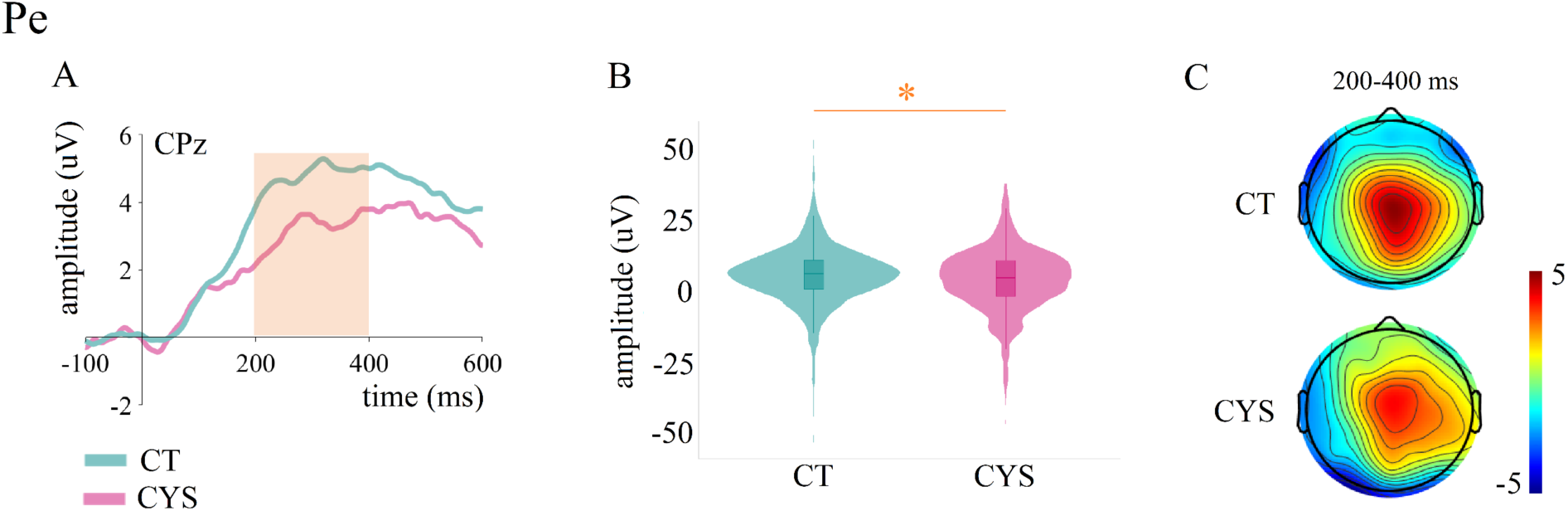
Averaged ERPs depicting error-related positivity (Pe) by group at CPz (A), plots showing distribution of amplitudes for Pe per group (trial-by-trial data) (B), and topographies per group between 200 and 400 ms (C). Shading in ERP plot indicates time window of interest.

### Correlations

Figure 5 shows all significant correlations with age (Panel A), between clinical scores and neural responses (Panel B), and between behavioral and neural responses (Panel C). Correlations’ direction and strength were identical between groups.

**Figure 5.**
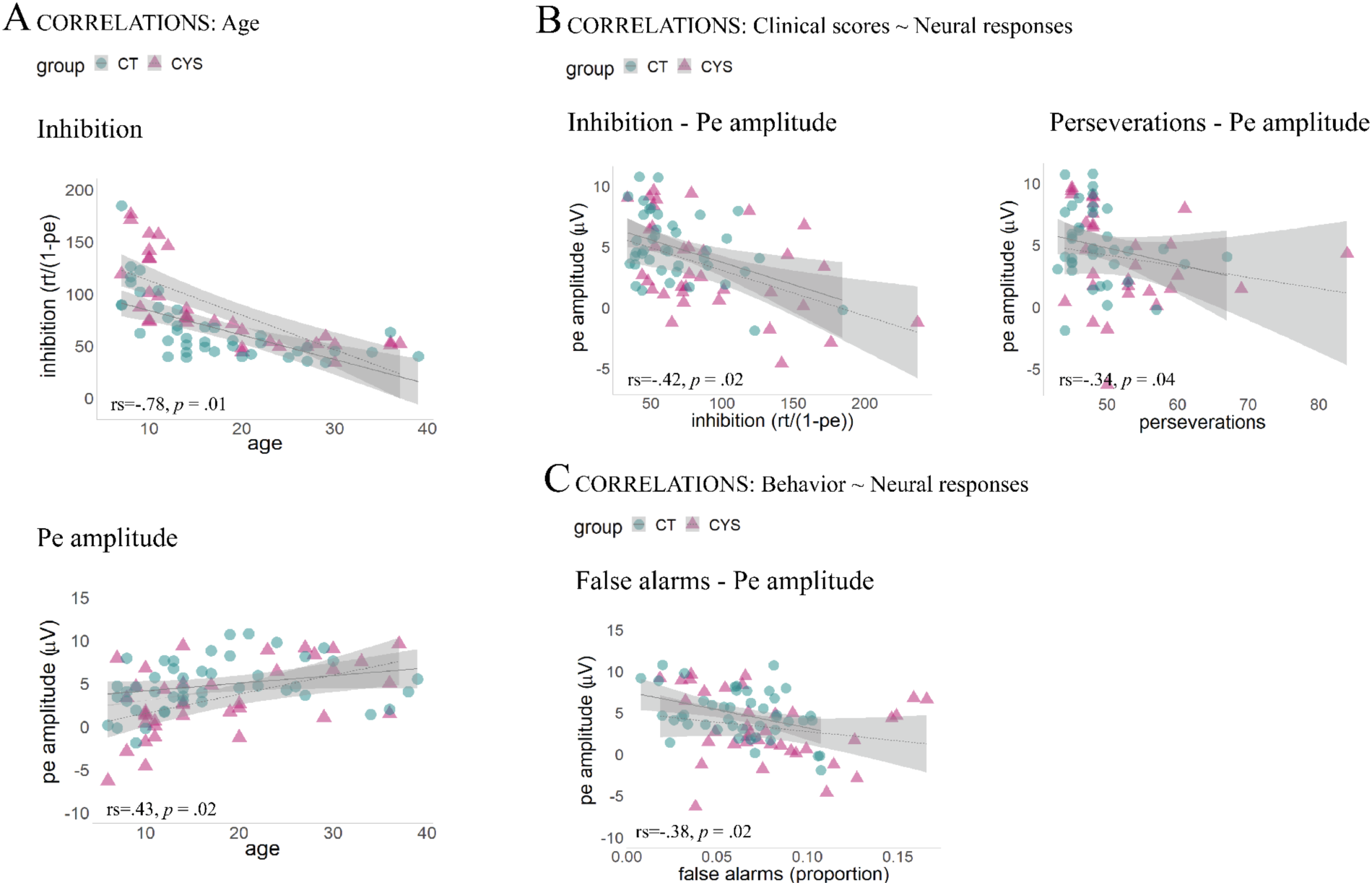
Spearman correlations between age and inhibition (Panel A); between clinical scores and neural responses (Panel B), and between behavioral and neural responses (Panel C).

Age correlated significantly with the inhibition composite measure (*r_s_*=-.78, *p* = .01), with older individuals from both groups performing better. Pe (*r_s_*=.43, *p* = .02) also correlated with age, with increased amplitudes in older individuals.

Only Pe amplitude correlated significantly with clinical scores. Associations between this component, inhibition composite score (*r_s_*=-.42, *p* = .02) and perseverations (*r_s_*=-.34, *p* = .04) were observed, with reduced Pe amplitudes relating to worse performance in those metrics. Pe amplitude was additionally associated with false alarm proportion: The higher the proportion of false alarms, the smaller the amplitude in the Pe time window (*r_s_*=-.38, *p* = .02).

## DISCUSSION

Utilizing standardized cognitive measures and EEG recordings, we investigated response inhibition and error monitoring in individuals with cystinosis.

### Inhibition

Neurocognitive assessments revealed that, when compared to their age-matched peers, individuals with cystinosis had greater difficulty inhibiting dominant and automatic verbal and motor responses. Similarly, a previous study investigating executive functioning in a group of children and adolescents with cystinosis reported differences in a set of D-KEFS tests, which included the Color-Word Interference Test (32), used here. In that particular task, approximately 30% of those with cystinosis scored below normal limits on the Color-Word Interference Test, suggesting marked difficulties in inhibitory processes in this population (32). An important finding is that cognitive function correlated with age in a similar fashion across groups. There is thus no evidence of cognitive decline with age in cystinosis, at least not one that deviates from what is expected in the general population.

During the EEG Go/No-Go task, and consistent with the performance on the standardized tests, the cystinosis group displayed poorer *d*-prime scores and longer reaction times when compared to the control group. While lower d-prime scores suggest diminished ability to adaptively balance demands to detect and respond vs inhibit responses, uniformly longer reaction times (whether for hits, false alarms, or post-false alarm trials) may be a consequence of the motor and processing slowing described in cystinosis (14, 32, 72). ERP analyses focused on components associated with different elements of response inhibition. In Go/No-Go tasks, the N2 is argued to index early, automatic inhibition (41–44) and/or conflict detection processes (45–47), whereas the P3 may be a marker of response inhibition (41, 48–52, 73), stimulus evaluation (53–55) and adaptive, effortful types of control (43, 44, 56). Given that ours and others’ behavioral findings suggest the presence of difficulties in response inhibition in cystinosis, reductions in ERPs indexing inhibitory processes were expected in this population.

As typically observed in the general population and shown here in controls, individuals with cystinosis had enhanced N2 and P3 amplitudes in response to correct rejections when compared to hits, suggesting that the inhibitory processes indexed by these components are overall preserved in this population. Moreover, no differences were found between the groups in the N2 time window, suggesting that early, more automatic inhibition might be maintained in cystinosis. In the P3 time window, however, individuals with cystinosis differed from their age-matched control peers, both in terms of mean amplitude and difference (correct rejections – hits) mean amplitude. Those with cystinosis presented larger P3 amplitudes and larger differences between correct rejections and hits when compared to controls. Though the N2 has been argued as a less reliable marker of response inhibition than the P3 (74–76), here, differences in the P3 time window between those with and without cystinosis could be mainly driven by significantly enhanced evoked responses to hits (see Figure 4A). One could thus argue that these differences in the P3 time window may not be indexing inhibitory difficulties per se but are the result of an overall increased response amplitude in cystinosis that is particularly prominent for hits, representing the majority of trials presented. Therefore, one should be cautious in interpreting the findings reported for the P3’s amplitudes and difference waves as they relate to the behavioral inhibitory difficulties observed in cystinosis. Interestingly, the amplitude enhancement in cystinosis amplitude appears to start much earlier, around 200 ms and to propagate steadily through time until around 700 ms after stimulus onset. We have previously reported increased P2 and P3a amplitudes in a sample of adults with cystinosis engaged in a passive auditory task (34) and argued that individuals with cystinosis may engage attention differently, which the current findings could be further evidence of. In future research, other processes potentially related to inhibition should be investigated in cystinosis. For instance, to be successful in a response inhibition task, one must maintain the task goal and the representation of the context or instance in which the response should be inhibited in working memory (77, 78). In other words, in this particular paradigm, one needs to remember the previous image in order to know which action to perform. Failure to do so at adequate levels of activation will probably result in marked difficulties in response inhibition processes (77, 79). Evidence regarding working memory difficulties in cystinosis is mixed (32, 34, 80), but our previous EEG work suggests difficulties in sensory memory in this population (34, 35), difficulties that could impact subsequent processing in working memory (81). Overall, a more detailed interrogation of the different components of response inhibition could be meaningful in clarifying the neural processes underlying the behavioral differences in inhibitory processing that we see here in cystinosis.

Lastly, one brief note regarding the P3 in the current data, which presents as a negative wave (see Figure 4A). Though atypical, the larger amplitudes in the younger participants and the long visual stimulus presentations may be contributing to the overall negativity of the response. Such pattern is observed in both groups and, therefore, we believe it does not impact the interpretation of the current findings.

### Error monitoring

Individuals with cystinosis presented reduced Pe amplitudes when compared to their age-matched peers. Pe reductions could suggest a weakened sense of error awareness (82). However, the post-error reaction times slowing in cystinosis in our data (reflected in longer reaction times in the trial after false alarm, Figure 2) suggests that those errors were registered, at least on a subset of error trials. Nevertheless, a possible explanation for the reduced Pe can be derived from the hypothesis that Pe may index a subjective or emotional error evaluation process, possibly modulated by the individual significance of the error (83). Individuals who commit errors more often (as those with cystinosis did in the present study), might attribute lower subjective or emotional significance to the errors made than those who rarely commit them. A lower attributed significance could thus result in a smaller Pe amplitude. This interpretation of Pe would further fit fMRI evidence suggesting that the rostral anterior cingulate, which is associated with affective processes, plays a key role in post-error processing (84, 85). Alternatively, reduced Pe amplitudes could be associated with slower processing speed, which has been reported in cystinosis. Previous studies have showed smaller Pe amplitudes for slow vs fast errors and in speed vs accuracy conditions. In those studies, larger Pe amplitudes were generally interpreted as reflecting more error evidence accumulation in a shorter period of time (80–83). Smaller Pe amplitudes in cystinosis could thus reflect the accumulation of less evidence and thus less certainty about the errors made. Interestingly, as can be seen in Figure S1 (Supplementary Materials), a small group of individuals with cystinosis presenting the most reduced Pe amplitudes, appear to also have slower reaction times. In cystinosis, difficulties in error monitoring could thus be, at least partially and for some individuals, explained by slower processing speed. Weakened working memory abilities could additionally explain why, though being slower in responding, some individuals with cystinosis made more false alarms than controls.

Lastly, that Pe amplitudes correlated, similarly in both groups, with number of commission errors, inhibition composite measure, and rate of false alarms, confirms this component’s functional association with and relevance to inhibitory and error monitoring processes.

### Study limitations

This study is not without limitations. First, our groups were not matched in terms of IQ, with the cystinosis group presenting significantly lower scores than the control group. Such differences could have impacted the results. Second, variables related to current health status (such as a measure of renal function) and compliance to treatment, which has been linked to better clinical outcomes (86), were not included in the present study but could be useful in understanding group- and individual-level differences. Relatedly, most individuals who participated in the study were relatively healthy and, thus, our sample may not be representative of the full spectrum of individuals living with cystinosis, which may impact generalization of findings. Third, whereas our data span a large age range that includes both children and adults, our participant numbers are not sufficient to power examination of the developmental trajectory of the inhibitory processes of interest. Such developmental changes would be expected based on well-characterized shifts in connectivity across the neural networks engaged by executive functions (87).

### Conclusions

In summary, this study provides the first EEG evidence of inhibition and error monitoring difficulties in cystinosis, and identifies processes important to consider in future research, such as attention, processing speed, and working memory. These findings have the potential to inform interventions to improve overall functioning in cystinosis. Relevant strategies may include a) repeating instructions of new information so that more information can be captured; b) teaching self-initiated “comprehension checking” strategies to help promoting independent management of working memory differences; c) providing a quiet, stable learning setting and reducing distractions in the environment that can tax or disrupt sustained attention; d) teaching and modeling self-regulation; e) practicing distractor blocking and attention switching; f) providing extended time for tests and assignments, to compensate for potential processing speed differences.

## ACKNOWLEDGEMENTS

We wish to thank Dr. Juliana Bates, who performed the clinical assessments, Elise Taverna and Danielle Newbury for their help with data collection and Dr. Frederick J. Kaskel for his help with recruitment. We extend our most sincere gratitude to the participants and their families for their interest, their involvement, and their time. This work was supported by a grant from the Cystinosis Research Network and a Eunice Kennedy Shriver National Institute of Child Health and Human Development U54 Grant (HD090260) to the Human Clinical Phenotyping Core of the Rose F. Kennedy Intellectual and Developmental Disabilities Research Center.

## COMPETING INTERESTS

The authors declare no conflicts of interest.

## AUTHOR CONTRIBUTIONS

AAF, SM, and JJF conceived the study. AAF, AB, and DJH collected and analyzed the data. AAF and AB wrote the first draft of the manuscript. SM, JJF, AB, and DJH provided editorial input to AAF on the subsequent drafts.

## Supplementary Materials

**Table S1.** Participants’ behavioral performance on the Go/No-Go EEG task: D-prime and reaction times

|  | controls | cystinosis |
| --- | --- | --- |
| **d-prime** | M=2.56; SD=0.37 | M=2.32; SD=0.42 |
| **hits (proportion)** | M=0.83; SD=0.06 | M=0.80; SD=0.06 |
| **false alarms (proportion)** | M=0.06; SD=0.03 | M=0.08; SD=0.04 |
| **RTs: hits** | M=396.91; SD=136.46 | M=428.24; SD=158.97 |
| **RTs: false alarms** | M=356.17; SD=160.74 | M=406.42; SD=201.51 |
| **RTs: after false alarms** | M=368.36; SD=176.83 | M=425.02; SD=205.23 |

**Figure S1.**
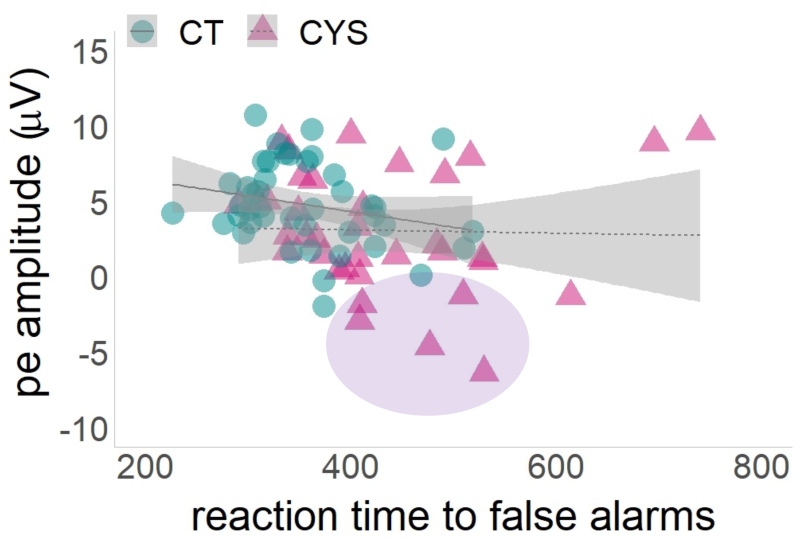
Spearman correlations between Pe amplitude and reaction time to false alarms.

## Notes

### Competing Interest Statement

The authors have declared no competing interest.

